# Single-nuclei RNA Sequencing Reveals Distinct Transcriptomic Signatures of Rat Dorsal Root Ganglia in a Chronic Discogenic Low Back Pain Model

**DOI:** 10.1101/2025.02.19.639130

**Authors:** Sydney M. Caparaso, Ishwarya Sankaranarayanan, David J. Lillyman, Theodore J. Price, Rebecca A. Wachs

## Abstract

Chronic low back pain (LBP), often correlated with intervertebral disc degeneration, is a leading source of disability worldwide yet remains poorly understood. Current treatments often fail to provide sustained relief, highlighting the need to better understand the mechanisms driving discogenic LBP. During disc degeneration, the extracellular matrix degrades, allowing nociceptive nerve fibers to innervate previously aneural disc regions. Persistent mechanical and inflammatory stimulation of nociceptors can induce plastic changes within dorsal root ganglia (DRG) neurons, characterized by altered gene expression, enhanced excitability, and lowered activation thresholds. Although these transcriptional changes have been described in other pain states, including osteoarthritis, they remain underexplored in discogenic LBP. To address this gap, this study represents the first application of comprehensive single-nuclei RNA sequencing of DRG neurons in a rat model of chronic discogenic LBP. Eighteen distinct DRG subpopulations were identified and mapped to existing mouse and cross-species atlases revealing strong similarities in neuronal populations with the mouse. Differential expression analysis revealed increased expression of pain-associated genes, including *Scn9a* and *Piezo2*, and neuroinflammatory mediators such as *Fstl1* and *Ngfr*, in LBP animals. Axial hypersensitivity, measured using grip strength, significantly correlated with increased expression of *Scn9a, Fstl1, and Ngfr,* which suggests their role in maintaining axial hypersensitivity in this model. These findings establish a relationship between DRG transcriptomic changes and axial hypersensitivity in a discogenic LBP model, identifying potential molecular targets for non-opioid treatments and advancing understanding of discogenic LBP mechanisms.

## 1. Introduction

Chronic low back pain (LBP) affects over 500 million people worldwide, resulting in significant disability and reduced quality of life [4,13,27,31]. Disc degeneration is highly associated with chronic LBP [11,15,42,44,62,70,73]. In up to 42% of chronic LBP patients, the pain is attributed to the disc, termed discogenic LBP [28,45,47]. However, existing treatments— including nonsteroidal anti-inflammatory drugs (NSAIDs) [60,71], opioids [32], and surgery [8]— often fail to provide long-term relief [18,43,59], highlighting the need to develop a comprehensive understanding of the mechanisms driving discogenic LBP.

Patients with discogenic LBP exhibit biological and anatomical changes within the disc. During disc degeneration, the extracellular matrix (ECM) breaks down, disrupting its ability to maintain hydration and native neuroinhibitory properties [33,36]. This breakdown allows nociceptive nerve fibers to innervate deep into the disc through axonal sprouting. Innervation has been observed in previously aneural regions including the nucleus pulposus (NP) core and the inner annulus fibrosus (AF) [7,20,24]. In addition, this mechanical breakdown of the disc is often observed with a catabolic microenvironment [10,34,74]. Quantitative sensory testing (QST) studies in LBP patients have shown decreased pain thresholds and heightened responses to mechanical [12,53] and cold stimuli [12,52] suggesting a process of nociceptor plasticity, or transcriptional changes to peripheral and central neurons with an implication on the dorsal root ganglia (DRG) as the primary site for sensory input. Importantly, studies of genetic trait loci in human DRGs have observed links between chronic pain conditions, such as temporomandibular disorder, and pathways involving nociceptor sensitization and immune responses [58]. However, there is a gap in understanding how transcriptional changes in nociceptors contribute to discogenic LBP.

Animal models have replicated several features of chronic discogenic LBP, including ECM breakdown, inflammation, and nociceptor innervation into the disc [2,40,49,55]. These animal models also recapitulate some subsets of pain-like behaviors like those exhibited in human patients [22,25,35,37,39,40,41,50,56]. Evidence from preclinical models of other pain states, such as knee osteoarthritis (OA), suggest that DRG neurons likely undergo plastic changes observed through the upregulation of ion channels and inflammatory markers [54,76]. Importantly, pain-related ion channel targeting has alleviated pain-like behavior in OA models; specifically, Nav1.7 [30], Trpv4 [68], and Piezo2 [54] have shown efficacy in mitigating OA progression and reducing pain-like response. Therefore, characterization of nociceptor plasticity in the context of chronic discogenic LBP could provide valuable insights into the molecular drivers of this disease.

Therefore, the goal of this work was to characterize DRGs in our discogenic rat LP model to understand nociceptive plastic changes in response to injury and correlation with pain-like behavior. To our knowledge, this study represents the first application of comprehensive single-nuclei RNA sequencing (snRNAseq) of rat DRG neurons. Herein, a high-resolution transcriptomic atlas of DRG subpopulations was generated to identify distinct nociceptor subtypes and their gene expression profiles in response to disc injury. In addition, our findings advance our understanding of DRG-specific changes associated with painful disc degeneration and highlight potential gene targets that correlate with pain-like behaviors.

## 2. Materials and methods

### 2.1. Study Design and Animals

A comprehensive dataset of dorsal root ganglia (DRGs) from 12 healthy adult female and male Sprague Dawley rats (Envigo) was deeply sequenced to establish a robust anchor dataset. This dataset was critical for accurately annotating rat neuronal subpopulations for subsequent pain studies. Herein, this study is referred to as the “anchor study,” as shown in (**Fig. 1**). For this anchor study, 6 rats (n=3 male and n=3 female) were harvested from both the University of Nebraska-Lincoln and the University of Texas at Dallas by two unique experimenters for a total of 12 rats. All procedures were conducted under approved Institutional Animal Care and Use Committee (IACUC) protocols at respective universities. This anchor dataset initially identified sixteen distinct clusters, providing unique DRG subpopulation annotations and establishing a foundational rat DRG atlas. After establishing the anchor dataset for snRNAseq of rat DRGs, a “chronic LBP study” examined transcriptional changes in DRG neurons in a model of chronic discogenic LBP as shown in (**Fig. 1**). For this study, 12 adult female Sprague Dawley rats (Envigo) from a larger cohort were randomly selected in (n=6 injured) and (n=6 sham) groups for sequencing. These samples were mapped onto the anchor dataset, enabling confident label transfer and precise classification of DRG subpopulations.

**Figure 1.**
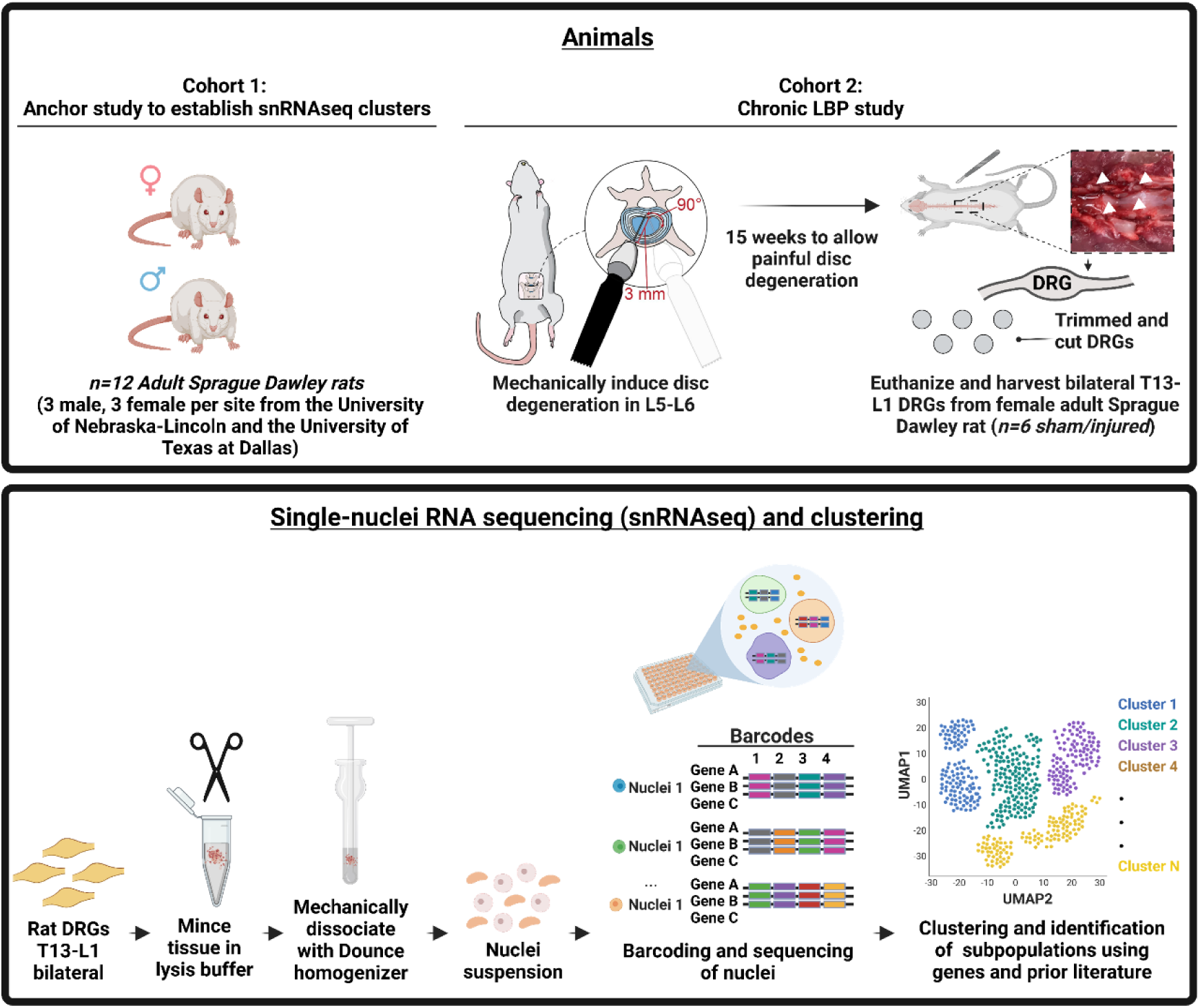
Schematic overview of experimental design for defining transcriptomic clusters in rat DRGs from chronic discogenic LBP model. **Top (Cohort 1)**: SnRNA seq of T13-L1 DRG was performed from uninjured control rats, and the analysis was conducted to create a high-resolution anchor dataset. T13-L1 bilateral DRGs from healthy male and female Sprague Dawley rats were harvested from UNL and UTDallas experimenters (n=12; 3 male and female per site at UNL and UTDallas). This dataset serves as a reference map for DRG subpopulations in healthy rats. **Top (Cohort 2)**: Adult female Sprague Dawley rats underwent surgical intervention to induce mechanical disc degeneration at the L5-L6 intervertebral disc level, followed by a 15-week period to allow for the development of painful disc degeneration. T13-L1 bilateral DRGs were harvested from sham and injured rats (n=6 per group). Labels defined from anchor study were transferred to the chronic LBP study. **Bottom**: Schematic of nuclei isolation for snRNAseq and clustering. Nuclei were isolated from DRGs, mechanically dissociated, and filtered to ensure single-nuclei suspension. Nuclei suspensions were then barcoded and sequenced using the Parse Bioscience. The data were analyzed for clustering visualization, revealing distinct clusters in the DRG. subpopulations based on gene expression profiles. Clusters were annotated to identify DRG subpopulations according to prior published human and mouse DRG datasets [6,61]. Dorsal root ganglia (DRG); low back pain (LBP); single-nuclei RNA sequencing (snRNAseq); University of Nebraska-Lincoln (UNL); University of Texas at Dallas (UTDallas).

### 2.2. Anchor study

#### 2.2.1. Humane euthanasia and DRG tissue harvest

Animals were humanely euthanized using CO_2_ inhalation as the primary method, followed by bilateral pneumothorax as a secondary method. DRGs from both sexes were carefully harvested based on methods from our lab [23]. In short, the spinous and transverse processes were removed, the spinal cord was carefully cut out, and the DRGs were extracted. During harvest, the connective tissue, spinal, and peripheral nerve processes were trimmed from the DRG explants. Bilateral T13-L1 DRGs were combined per animal for snRNAseq and flash frozen in liquid nitrogen before further processing. These DRG levels were selected as L1 DRGs are a primary contributor to innervation in the L5-L6 disc in rats [49,55,65].

##### 2.2.1.1. DRG nuclei isolation from adult Sprague Dawley rats

Prior to handling DRG samples, all surfaces and tools were cleaned with 70% ethanol and an RNase decontamination solution (RNAzap, Thermofisher). Nuclei were isolated based on methods from the manufacturer’s protocol (Parse Evercode V2 kit) and literature [6]. In short, T13-L1 bilateral DRGs per animal were chopped into <1mm small pieces using Bonn scissors (14084-08, FST) in a 1.5 mL DNA LoBind tube (Eppendorf, 0030108418) with 500 µL of homogenization buffer followed by mechanically dissociating using the Dounce homogenizer with 15 pestle strokes (**Fig. 1**). The buffer was freshly prepared by combining 0.25 M sucrose, 150 mM KCl, 5 mM MgCl₂, 1 M Tris buffer, pH 8.0, 0.1 mM DTT (ThermoFisher, D1532), cOmplete protease inhibitor (Sigma, 11836153001), 0.1% Triton X-100 (Sigma, T8787), and 0.2 U/µL RNAsin (Promega, N2515). Then, the tissue was filtered through a 70 µm cell strainer (Corning, 352360) into a 50 mL conical tube. The nuclei suspension was centrifuged at 500 x g for 5 minutes at 4°C, and the supernatant was discarded. The nuclei pellet was washed once in wash buffer containing 1% RNase-free BSA, 0.2 U/µL RNase inhibitor, 1X DPBS without calcium and magnesium (ThermoFisher, 14190144) and centrifuged at 500 x g for 5 minutes. The nuclei were counted using 0.4% Trypan Blue and hemocytometer to determine the number of cells. (ThermoFisher, 15-250-061) and hemocytometer. Nuclei were fixed and processed using the manufacturer’s instructions using the Evercode Nuclei Fixation v2 kit (Parse Biosciences, ECF2003). The libraries were prepared according to the manufacturers’ instructions (Parse Evercode V2). Libraries were sequenced using Nextseq2000 at the University of Texas at Dallas Genome Core.

##### 2.2.1.2. Bioinformatics analysis

The sequenced libraries were processed using Parse Biosciences Evercode Software Suite and adapted from prior methods [6]. Initial quality control steps included filtering out nuclei with low gene counts (<200 genes), high mitochondrial RNA content (>10%), or suspected doublets to retain only high-quality nuclei, were processed using the Seurat pipeline. The resulting expression profiles were normalized and scaled to account for differences in sequencing depth. Dimensionality reduction was performed using clustering with Uniform Manifold Approximation and Projection (UMAP) for clustering and visualization of DRG subpopulations. To annotate the anchor study dataset, a prior published single-cell cross-species atlas from Bhuiyan et al. [6] was used to transfer levels using Conos. This established a detailed anchor set for rat DRG populations which was subsequently used as a reference to assign labels to a dataset containing both sham and injured animals. Differential expression analysis was conducted using the DEseq2 package within the labeled clusters between sham and injured animals. Data visualization was conducted in R. Data can be accessed at https://paincenter.utdallas.edu/sensoryomics/.

### 2.3. Chronic low back pain study

#### 2.3.1. Animal acclimation and husbandry

All animal procedures followed the National Institutes of Health guidelines and the Public Health Service Policy on Humane Care and Use of Laboratory Animals and were approved by the IACUC at the University of Nebraska-Lincoln. Adult female Sprague Dawley rats (Envigo) were housed under a reverse 12-hour light/dark cycle with ad libitum access to food and water. Before the start of the study, each animal was acclimated to experimenters for a minimum of five minutes per day for eight days.

#### 2.3.2. Surgical procedure to elicit chronic low back pain

During surgery, rats were anesthetized using 2-3% isoflurane in oxygen and given a single subcutaneous injection of Buprenorphine SR (0.75 mg/kg) for post-operative pain management. The L5-L6 disc was located using the iliac crest as a landmark, and tissue dissection was performed according to previously established methods in our lab [40]. The abdomen was shaved and disinfected with betadine and iodine. A midline incision was made along the ventral abdomen at the level of the iliac crest, extending through the skin, subcutaneous tissue, and fascia to expose the retroperitoneal space while avoiding major blood vessels. The L5-L6 disc was bilaterally punctured using a 3 mm dissecting needle (Roboz, RS-6066) and swept in a 90° arc six times in the transverse plane, following an established protocol [40]. The abdominal incision was closed using a continuous subcuticular suture pattern. Animals were observed 1 hour after surgery and then every 12 hours for 3 days to monitor well-being and wound healing. Post-operative behavioral assessments began after a two-week recovery period. By the end of the study two sham animals were euthanized due to surgical complications or illness, making the final animal numbers per group: sham = 10, injured = 12. These animals made up the final pool from which animals were randomly selected for snRNAseq.

#### 2.3.3. Animal health pain-like behavior assessments

Prior to all behavioral testing, animals were acclimated to equipment for at least five minutes per animal per apparatus. Animals were randomly assigned to groups and experimenters were blinded to all animal assignments. Animals were housed under a reverse light cycle and all behavioral data was collected with red light to reduce stress. Health and well-being were monitored bi-weekly by measuring weight, and any weight loss exceeding 10% from the previous measurement prompted veterinary consultation. Pain-like behaviors were evaluated bi-weekly post-operatively using two evoked assays (grip strength and pressure algometry). Prior to the start of the study, two baseline collections per animal were averaged for each assay.

##### 2.3.3.1. Grip strength

Axial hypersensitivity was quantified using grip strength (San Diego Instruments, 2325-0063) following methods previously described by our lab [40]. Animals underwent acclimation to the testing environment for at least 5 minutes before each test. In short, animals were held by the hindquarters which allowed them to grip a mesh metal wire that was connected to a grip strength force sensor. After the animal gripped the bar, the animal hold was then shifted to the base of the tail and the animal was gently and uniformly pulled back from the bar over the course of 1-1.5 seconds until it released the mesh. Three trials were recorded, and the average maximum force (N) was used. Grip strength values were normalized to baseline measurements for each animal.

##### 2.3.3.2. Pressure algometry

Pressure sensitivity at the L5-L6 region was assessed using an electronic von Frey aesthesiometer (IITC, 2391) using a blunt probe following methods previously described by our lab [40]. In short, prior to testing, each animal was acclimated to the testing environment for at least 5 minutes. During testing, the animal was carefully wrapped in a clean cotton t-shirt and loosely restrained by one experimenter, while a second experiment applied the blunt probe to the skin over the L5-L6 area. The pressure gradually increased until the animal demonstrated a pain-like response, such as avoidance behavior or vocalization. Two measurements were taken for each animal, and the average was recorded. All pressure thresholds were normalized to the baseline measurements for each animal.

#### 2.3.4. Motion segment processing

Tissue was processed following methods previously described by our lab [40]. In short, the L5-L6 motion segments were carefully isolated from each animal, cleaned, and fixed in 3 mL of 4% paraformaldehyde (PFA, Sigma Aldrich, 441244-1KG) for 24 hours at room temperature, with continuous agitation at 180 rpm on an orbital shaker. After fixation, the motion segments were rinsed three times for 15 minutes each with 1X PBS and subsequently placed in 3 mL of Immunocal (Fisher Scientific, NC9044643) to decalcify for 18 hours at room temperature with constant agitation on an orbital shaker. Once decalcified, the motion segments were washed three times with 1X PBS for 15 minutes each and then soaked in a 30% sucrose solution (wt/v) prepared in 1X PBS for 24 hours at 4°C. The segments were cryoembedded in Optimal Cutting Temperature Compound (Scigen 4586), frozen at −80°C, and sectioned in the sagittal plane into 40 μM slices. One injured animal was excluded from analysis due to issues with tissue harvesting, making the final animal numbers for subsequent motion segment histological analyses and behavioral analyses: sham = 10, injured = 11.

#### 2.3.5. Immunohistochemistry

Immunohistochemistry (IHC) was performed to assess the degree of nerve innervation in the disc. A peptidergic C-fiber marker for calcitonin gene-related peptide (CGRP) was selected due to prior demonstrated efficacy of staining nerve fibers in the disc [2,40,48,57]. Three 40 μM thick sections of the L5-L6 motion segments from each animal were post-fixed in 4% paraformaldehyde (PFA) for 15 minutes, followed by two 5-minute washes in 1X PBS and a final 5-minute wash in 1X PBST. The sections were then blocked for 1 hour in a blocking buffer consisting of 1X PBST, 3% goat serum (Sigma-Aldrich, G9023), and 0.01% Tween-20 (Fisher Scientific, BP337-100). Following blocking, the sections were then incubated overnight (16 hours) with a primary antibody diluted in blocking buffer, using 1:100 mouse anti-CGRP (Abcam, Ab81887). After incubation, the slides were washed three times for 15 minutes each with PBST, then incubated for 2 hours with a secondary antibody 1:500 goat anti-mouse AF488 (Abcam, Ab150117) diluted in blocking buffer. Following secondary incubation, the sections were washed three additional times with PBST for 15 minutes each, followed by a 10-minute incubation with 1:1000 DAPI (ThermoFisher, D1306) in 1X PBS for nuclear staining. Finally, the sections were mounted using Prolong Gold antifade reagent (ThermoFisher, P36930). Slides were imaged using a Zeiss LSM 800 confocal microscope (Carl Zeiss Microscopy) with a 10X objective. Whole motion segments were captured using the tiling feature and z-stacks to visualize the entire section thickness. Both brightfield and fluorescent images were taken with the following excitation/emission settings: 353/465 nm for DAPI and 493/517 nm for AF488.

#### 2.3.6. Image analysis

Immunohistochemistry images were first processed using ZEN Blue (Carl Zeiss Microscopy), where Z-stack projections were generated and saved as .czi files. These files were then imported into QuPath-0.4.3 (Pete Bankhead) for further analysis and annotation of disc regions. The wand and brush tools were used to create annotations on each section, representing distinct disc regions, including the dorsal ligament, dorsal annulus fibrosus (AF), nucleus pulposus (NP), the inner two-thirds of the ventral AF, the outer one-third of the ventral AF, granulation tissue, and ventral ligament. After completing the annotations, the ‘Pixel Classifier’ tool was applied to threshold CGRP-positive staining within each region. The percentage of positive staining was calculated by normalizing the CGRP-positive area to the total area of the respective disc region.

#### 2.3.7. Statistical analysis

Behavioral data are presented as mean ± 95% confidence interval (CI). IHC data are presented as mean ± standard deviation. All data were analyzed using GraphPad Prism V9. Normality was assessed using a Shapiro-Wilk test to determine subsequent analyses. Nerve IHC were analyzed using a Kruskal-Wallis test with Dunn’s post hoc. Behavioral data were analyzed using a Two-Way ANOVA with Dunnett’s post hoc test. Results were considered statistically significant when p≤0.05.

Differential gene expression analysis was performed between sham and injured animals, looking at genes averaged across all neuronal clusters and within specific clusters. Pseudobulked count matrices were generated by summing raw counts per gene across all neuronal nuclei within each sample before applying DESeq2 normalization and modeling. Preprocessing included quality filtering and normalization to account for variations in sequencing depth. Genes were considered significantly upregulated in injured animals with a Log Fold Change (LogFC) threshold of 1.35, and p-adjusted ≤0.05. Differential expression analysis between sham and injured animals within each defined cluster was conducted using a non-parametric Wilcoxon rank-sum test. Comparisons between select genes across neuronal clusters were conducted using a non-parametric Mann-Whitney U test. Spearman correlations were calculated between gene expression levels and pain-like behavior metrics (grip strength and pressure algometry) at week 15 as well as CGRP-positive nerve staining in the disc.

## 3. Results

### 3.1. Anchor study

#### 3.1.1. snRNAseq and clustering

The anchor set consisted of 13,346 nuclei, which were clustered into 16 distinct clusters (**Fig. 2A**). High-resolution snRNA-seq metrics demonstrated robust sequencing quality, with a median of 1,830 transcripts per cell, a median of 1,155 genes per cell, and a mean of 97,967 reads per cell (**Fig. 2B-C**). The average mitochondrial RNA content across samples remained below 10%, suggesting minimal cell damage or lysis during processing. These metrics indicate the data had sufficient sequencing depth and quality for downstream analysis. Neuron-specific genes were examined, and strong expressions of both *Snap25*, and *Rbfox3* (NeuN) were observed across neuronal clusters, with lower expression in Schwann cell cluster 2, which expresses Schwann cell markers (**Fig. 2D, E**). Dot plot analyses revealed the percentage of cells expressing various genes across clusters, highlighting specific expression patterns. *Ntrk3* was highly expressed in clusters 4 and 13, *Trpm8* was predominantly expressed in cluster 15, and *Il31ra* showed strong expression in cluster 14, along with their corresponding average expression levels (**Fig. 2F**). UMAP projections highlighted spatial distribution and distinct expression patterns of genes such as *Calca, Il31ra, Ntrk2,* and *Gfra2* in specific cluster populations (**Fig. 2G**). The clusters were confidently annotated using our previously published dataset, which includes genes from mouse species [6].

**Figure 2.**
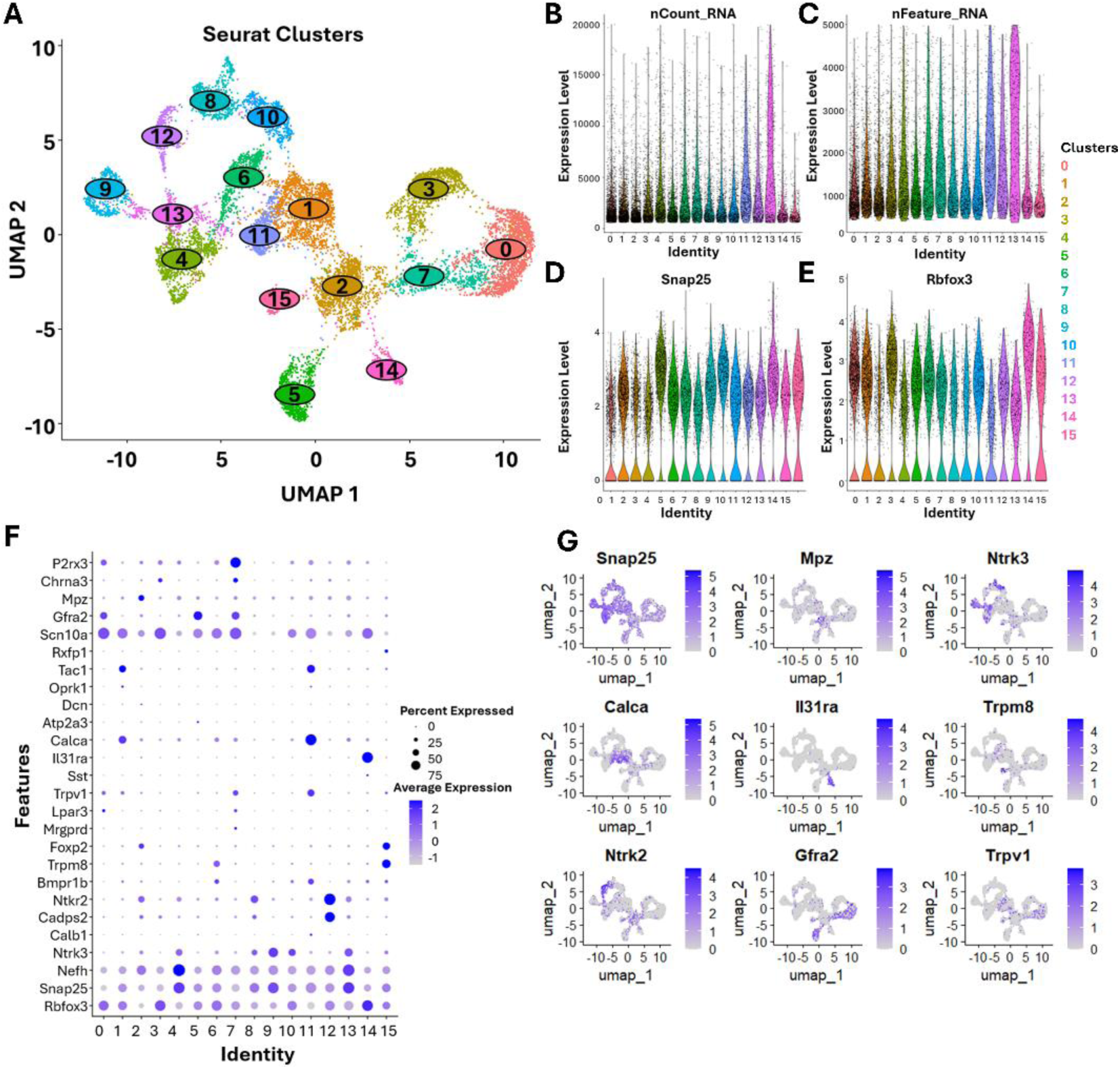
DRG sequencing and clustering analysis in the anchor study to establish subpopulation labels. (A) UMAP plot of predicted clusters for T13-L1 bilateral DRGs harvested from male and female Sprague Dawley rats (n=6 male and female). Quality control metrics for RNA sequencing in the anchor dataset: (B) Volin Plot showing total number of molecules detected within a cell (nCount_RNA) and (C) Volin Plot showing number of genes detected in each cell (nFeature_RNA). (D) Volin Plot showing the expression levels of the neuronal marker *Snap25* and (E) *Rbfox3* (neuronal nuclei) across clusters. (F) Dot plots showing the percentage of cells expressing key genes across clusters. UMAP showing gene expression markers (G). Uniform Manifold Approximation and Projection; (UMAP). N=6 rats per group.

Using Conos label transfer, eighteen neuronal subtypes were identified across 13,346 nuclei (**Fig. 3A**). These annotations were based on the harmonized atlas, of which 15 neuronal subtypes have been described as having at least one known function in rodent models [6]. UMAP projections revealed distinct clustering of nuclei, with populations showing similar proportions in both mouse and rat DRG datasets (**Fig. 3B-C**). The subpopulations include A-fiber subtypes such as Pvalb-expressing proprioceptors, Ntrk3high+Ntrk2 Aβ-rapid-adapting (RA) low-threshold mechanoreceptors (LTMRs), Ntrk3high+S100a16 Aβ-slow-adapting (SA) LTMRs, Ntrk3low+Ntrk2 Aδ-LTMRs, Calca+Bmpr1b Aδ-high-threshold mechanoreceptors (HTMRs), and Calca+Smr2 Aδ-HTMRs. The C-fiber subtypes include Calca+Sstr2, Calca+Adra2a, and Calca+Dcn nociceptors; Calca+Oprk1, a nociceptor expressing the opioid receptor kappa 1; Trpm8 cold nociceptors; and Th C-LTMRs. Additional C-fiber subtypes include Mrgprd nociceptors, Mrgpra3+Mrgprb4 C-LTMRs, Mrgpra3+Trpv1 pruriceptors, and Sst pruriceptors. Notably, Rxfp1, a rare subtype previously described to express high levels of Trpv1 [6], was also identified in our anchor dataset. Lastly, a small representation of an Atf3 cluster was also identified, characterized by the expression of this neuronal injury-associated gene. While the clusters represent shared cell types across species, specific clusters exhibited unique distributions, as shown in the pie chart, reflecting species-specific transcription patterns (**Fig. 3C-D**). Notably, the proportion of Th C-LTMRs was higher in mice (13.37%) compared to rats (8.68%), whereas Pvalb-expressing proprioceptors were more prevalent in rats (7.96%) than in mice (4.06%), but these differences may also be due to the level of DRGs assessed. In addition, the proportion of Ntrk3high+Ntrk2 Aβ-RA LTMRs was higher in rats (4.88%) compared to mice (2.81%). Dot plot analyses comparing the anchor dataset and the reference dataset demonstrated the expression profiles of key genes across clusters in both species (**Fig. 3D-E**). *Trpm8*, associated with cold sensing, was predominantly expressed in specific clusters in both mouse and rat, showing similar gene expression patterns across species. Pvalb, a marker of proprioceptive neurons, exhibited consistent cluster enrichment patterns alongside genes such as *Esrng*, *Vsnl1*, and *Tafa2* in both species. Similarly, the peptidergic cluster population Calca+Bmpr1b displayed unique clustering patterns, characterized by distinct expression of genes including *Tafa1*, *Kcnq5*, and *Csmd3*. These findings highlight both conserved and species-specific transcriptional features in neuronal subtypes. Next, using a mouse reference dataset, Bhuiyan et al. [6] and https://paincenter.utdallas.edu/sensoryomics/sensoryomics-dashboard/ [63], we identified 49 rat-specific genes distinguishing rat-specific transcriptional features. These genes were present in this rat dataset but were absent from the mouse reference dataset (**Fig. 3F**). These genes included *Abcg3l1* (ATP-binding cassette, subfamily G)*, Dennd2b* (DENN domain containing 2B)*, Dsp* (Desmoplakin) a gene involved in cell adhesion*, Kcne2* (Potassium Channel, Voltage-Gated Subfamily E Regulatory Beta Subunit 2)*, Lhfpl6* (Tetraspan subfamily member 6)*, Nrdc* (Nardilysin), a metallopeptidase that regulates growth factors and inflammatory response, and *Tlcd4* (TLC Domain-Containing 4), which were not detected in the mouse dataset [6].

**Figure 3.**
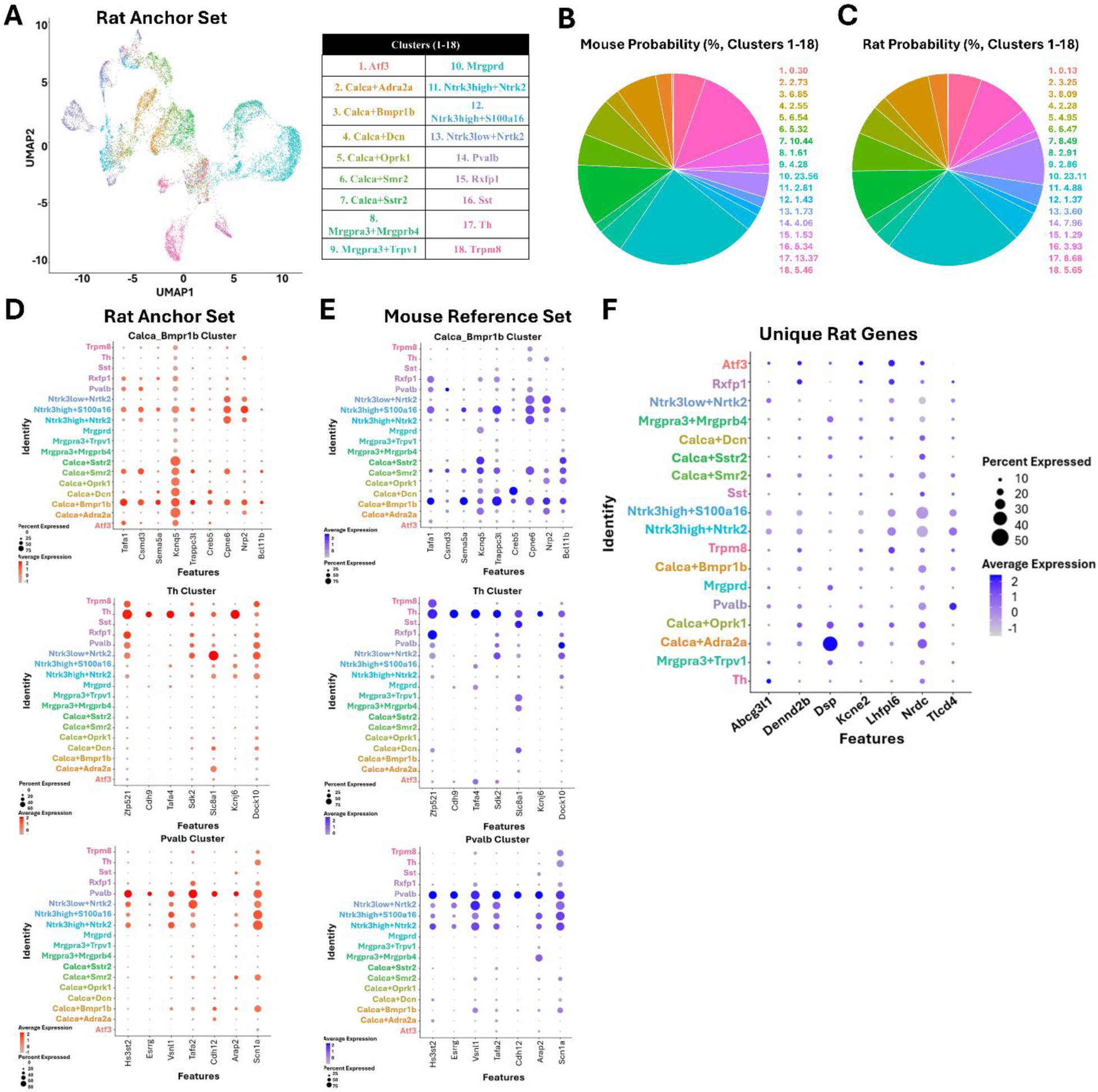
Species-Specific and Conserved Neuronal Subtypes in DRG. (**A**) UMAP showing 18 neuronal subtypes after Conos label transfer with labelled clusters of neuronal subtypes. (**B-C)** Species-specific differences in neuronal subtype proportions between mouse and rat. (**D-E**) Dot plot analyses demonstrated conserved gene expression profiles across mouse and rat for clusters. (**F**) Comparative analyses identified 49 rat-specific genes absent in mouse datasets, including *Abcg3l1, Dennd2b, Dsp*, and *Kcne2*, which are associated with cell adhesion, structural integrity, and inflammatory responses. Six rats (n=3 male and n=3 female) were harvested from both the University of Nebraska-Lincoln and the University of Texas at Dallas by two unique experimenters for a total of 12 rats. Uniform Manifold Approximation and Projection; (UMAP).

### 3.2. Chronic LBP study

#### 3.2.1. snRNAseq and clustering

Libraries were generated from the bilateral T13-L1 dorsal root ganglia (DRG) of a chronic low back pain cohort, which included 12 female rats—6 from the injured group and six from the sham group. A total of 39,058 nuclei were recovered across both conditions. Nuclei had a median of 1,162 transcripts, and the sequencing depth was an average of 16,395 reads per nuclei. Conos label transfer was applied to the chronic lower back pain cohort using the previously annotated anchor set. UMAP projections again identified 18 neuronal subpopulations. The cluster subtypes showed broad similarities between the injured and sham groups (**Fig. 4A-B**). Cluster proportions between sham and injured rats were compared (**Fig. 4C**). While most clusters were consistently represented in both groups, we observed slight increases in the proportions of injury-associated clusters in the injured animals. For example, the Ntrk3low+Ntrk2 (Aδ-LTMRs) cluster showed a cell proportion of 3.2% in the injured group compared to 2.3% in the sham group, reflecting injury-induced transcriptional changes.

**Figure 4.**
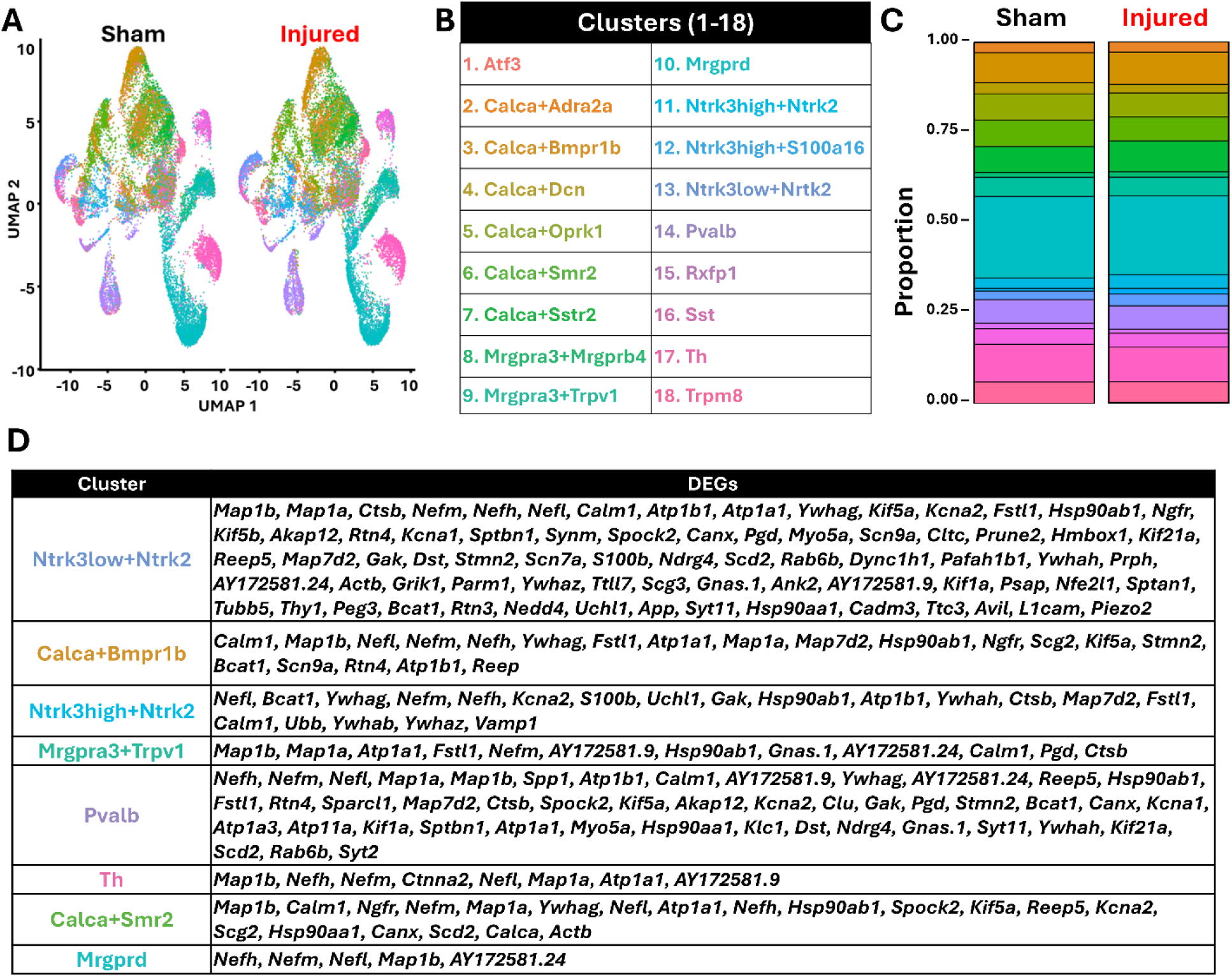
snRNAseq atlas comparing sham and injured DRG subpopulations in a chronic LBP model. (**A**) UMAP projection showing 18 distinct clusters identified from the anchor dataset; labels were transferred to the chronic LBP dataset and consistent across sham and injured animals. (**B**) Clusters annotated using the Bhuiyan et al. [6] dataset for both anchor and chronic LBP datasets, showing 18 defined clusters. (**C**) Relative proportions of each DRG subpopulation in sham and injured animals. (**D**) All differentially expressed genes by cluster. N=6 animals per group. Uniform Manifold Approximation and Projection; (UMAP).

Differential gene expression analysis was performed between sham and injured animals across all neuronal clusters as well as within specific clusters. DEGs were identified between sham and injured animals across clusters (**Fig. 4D**). This approach inherently dilutes cluster-specific signals, as variations in gene expression within specific subpopulations may be masked by the larger, heterogeneous population. The highest proportion of DEGs was observed in the Ntrk3low+Ntrk2 (Aδ-LTMRs), Calca+Bmpr1b (Aδ-HTMRs), and Pvalb (A-fiber proprioceptors) populations.

In the injured group, several pain-associated genes showed significant upregulation compared to sham log2FC of 0.44 or higher. For instance, within the Ntrk3low+Ntrk2 population (Aδ-LTMRs) cluster, *Scn9a* and *Piezo2*, key ion channels involved in pain sensitivity, were significantly upregulated in injured animals. Specifically, *Scn9a* exhibited a log2 fold change of 0.58 while *Piezo2* showed a log2 fold change of 0.44. The Ntrk3low+Ntrk2 population (Aδ-LTMRs) are known to innervate hair follicles and detect hair deflection [1] but have no known role in disc innervation. Additionally, *Ngfr* (nerve growth factor receptor), a marker linked to neurotrophic signaling in pain pathways, displayed significantly increased expression within this cluster in injured animals with a log2FC of 0.66. Other genes associated with disc degeneration and cellular senescence were also significantly elevated in the injured group in this cluster. Importantly, *Spock2* (sparc/osteonectin, cwcv and kazal-like domains proteoglycan 2), related to extracellular matrix integrity and implicated in disc degeneration and *Fstl1*, related to nucleus pulposus cell senescence and intervertebral disc degeneration were significantly increased in the Ntrk3low+Ntrk2 population (Aδ-LTMRs) (**Fig 4D**). These data suggest injured animals may exhibit alterations in genes associated with pain signaling and disc degeneration mechanisms. The highest proportion of differentially expressed genes were found in the Ntrk3low+Ntrk2 (Aδ-LTMRs), Calca+Bmpr1b (Aδ-HTMRs), and Pvalb (A-fiber proprioceptors) populations (**Fig. 4D**). Consistent DRG clusters across sham and injured animals suggest that injury does not alter overall DRG subpopulations but affects gene expression within existing neuronal subtypes.

Differential expression analysis was conducted across all neuronal clusters to identify transcriptional changes following injury. A total of 75 representative DEGs with an adjusted p-value (p-adj) < 0.05 are shown (**Fig. 5A**), selected from a larger set of significant DEGs identified across all neuronal clusters. These genes were categorized into functional groups, including neuronal cytoskeleton and axonal transport, ion channels and transporters, chaperones and heat shock proteins, signal transduction and synaptic proteins, metabolism and enzymes, extracellular matrix and adhesion, and other. Expression patterns of selected genes of interest are shown across neuronal clusters (**Fig. 5B–E**). *Ngfr* expression was broadly increased in injured animals, particularly in Pvalb (A-fiber proprioceptors) and Ntrk3high+S100a16 (Aβ-SA LTMRs) clusters (**Fig. 5B**). *Scn9a* expression was elevated following injury, particularly in Trpm8 (cold nociceptors) and Mrgpr3+Mrgprb4 (C-LTMRs) clusters (**Fig. 5C**). *Fstl1*, an extracellular matrix-related gene, exhibited cluster-specific injury-induced upregulation, with the highest expression observed in Pvalb (A-fiber proprioceptors) and Mrgpr3+Mrgprb4 (C-LTMRs) clusters (**Fig. 5D**). Similarly, Spock2 showed distinct cluster-dependent changes, with the most pronounced increases in Pvalb (A-fiber proprioceptors) and Ntrk3low+Ntrk2 (Aδ-LTMRs) clusters (**Fig. 5E**). These data suggest injury-induced transcriptional adaptations across neuronal subtypes, highlighting potential mechanisms contributing to altered neuronal function and extracellular matrix remodeling in response to injury.

**Figure 5.**
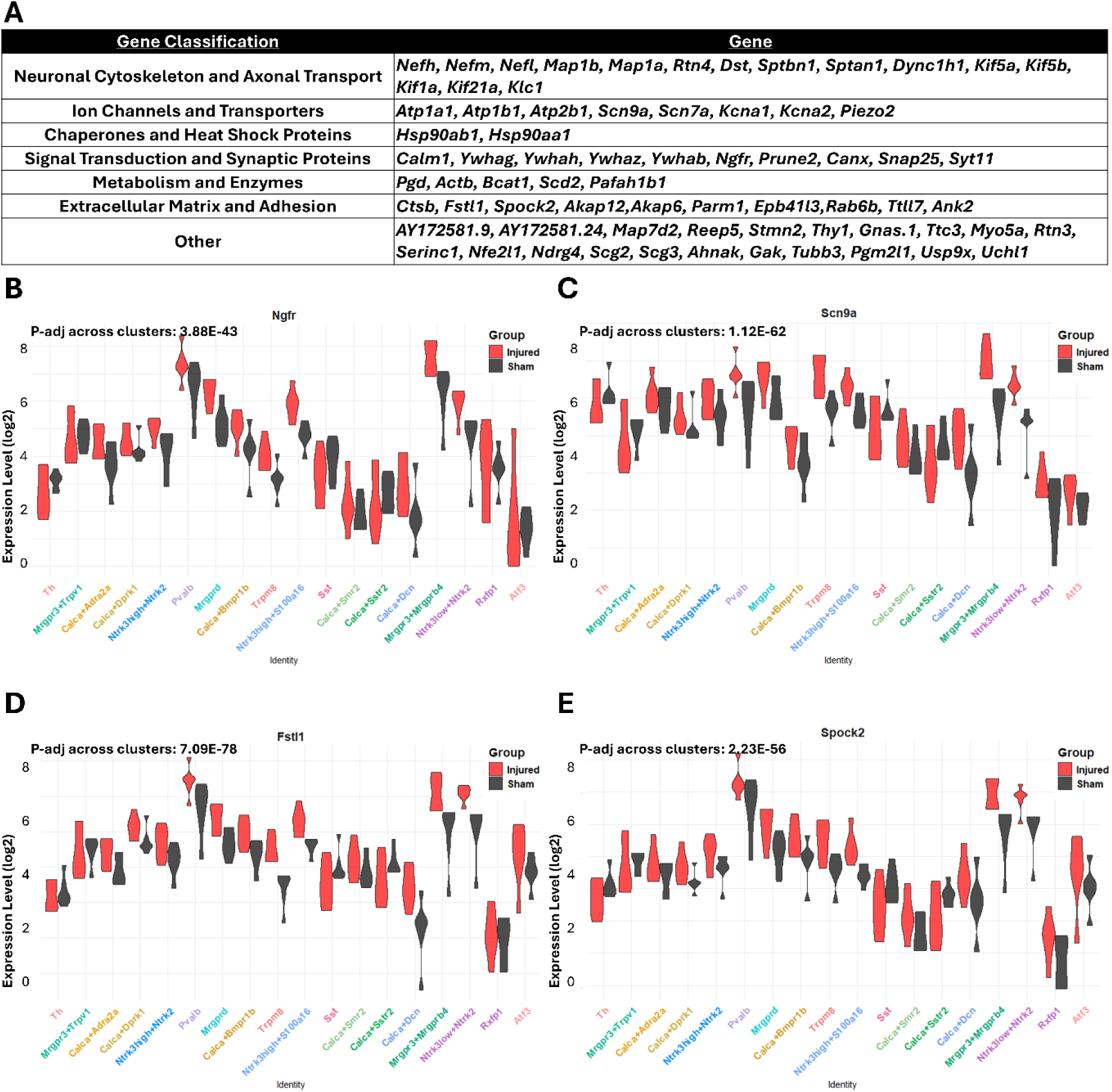
Differential expression of key genes across neuronal clusters in sham and injured animals. **(A)** Gene classification table grouping differentially expressed genes (p-adj <0.05) by functional category. Showing 75 DEGs from a larger pool. **(B-E)** Violin plots of *Ngfr* **(B)**, *Scn9a* **(C)**, *Fstl1* **(D)**, and *Spock2* **(E)** expression across neuronal clusters, comparing sham (black) and injured (red) animals. Expression levels are log-transformed (log2) for visualization. N=6 animals. P-adj value across clusters per gene displayed on each violin plot.

#### 3.2.2. Differentially expressed pain-related genes correlate with pain-like behavior

Animals developed progressive axial hypersensitivity, emerging around week 8 and remaining statistically significant compared to sham controls through week 15, with injured animals exhibiting grip strength reduced to ≤82% of baseline (**Fig. 6A and B**). Significant correlations were identified between grip strength and expression levels of *Scn9a* (p=0.0102), *Ngfr* (p=0.0275), *Fslt1* (p=0.0025), and *Spock2* (p=0.0205) across all clusters, suggesting these genes may play a role in the maintenance of axial hypersensitivity. A trend was also observed between grip strength and *Cdkn2b* (p=0.067) (**Fig. 6C**) across all clusters.

**Figure 6.**
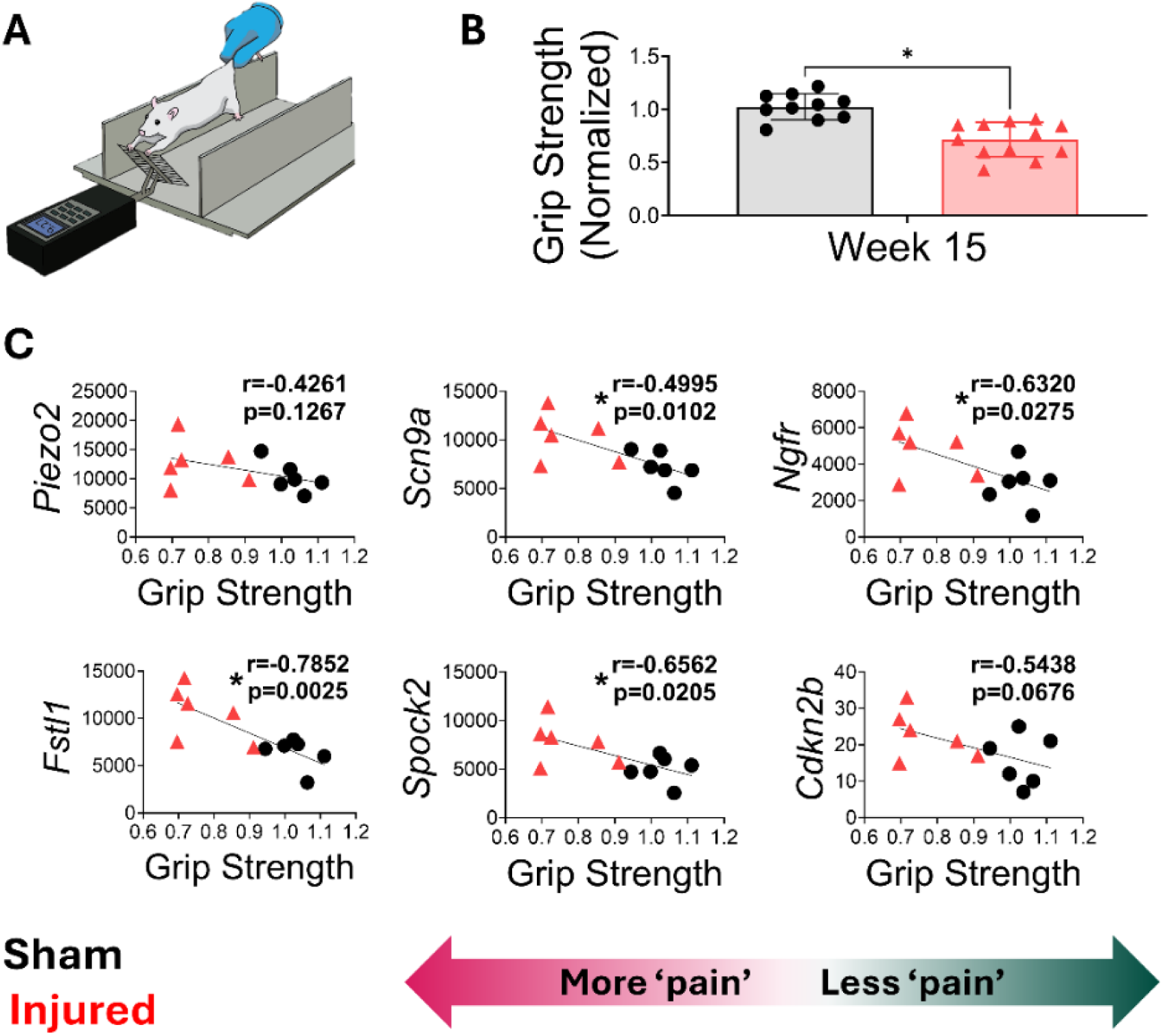
Spearman correlations of differentially expressed genes with axial hypersensitivity in sham and injured animals measured by grip strength. **(A)** Schematic of the grip strength assay to evaluate axial hypersensitivity. **(B)** Normalized grip strength at week 15 post-injury, showing significantly lower grip strength in injured animals indicating axial hypersensitivity. **(C)** Spearman correlations between differentially expressed genes relevant to pain-like behavior and disc degeneration (*Scn9a, Ngfr, Fstl1, Spock2, and Cdkn2b*) and grip strength. Significant correlations were found for *Scn9a*, *Ngfr*, *Fstl1*, and *Spock2*, suggesting these genes may contribute to persistent axial hypersensitivity. A trend toward significance was observed for *Cdkn2b* (p=0.067). N=6 animals per group. Black symbols represent sham, red symbols represent injured animals. *=p≤0.05. The arrow indicates that higher grip strength values correspond to less pain-like behavior, reflecting a value closer to the baseline (1).

Significant differences in pressure algometry were observed between sham and injured animals from weeks 8 to 15, with injured animals exhibiting pressure thresholds reduced to ≤81% of baseline (**Fig 7A-B**); however, no significant correlations or trends were observed between pressure algometry and differentially expressed genes across all clusters (**Fig. 7C**). Together, these data suggest that axial hypersensitivity is a prominent pain phenotype in this discogenic LBP model with correlations to pain-related genes in the DRG. In addition, localized touch sensitivity, measured through pressure algometry, showed a significant decrease in injured animals suggesting a pain phenotype present in our animals; however, no significant correlations with DRG changes were observed with pressure algometry.

**Figure 7.**
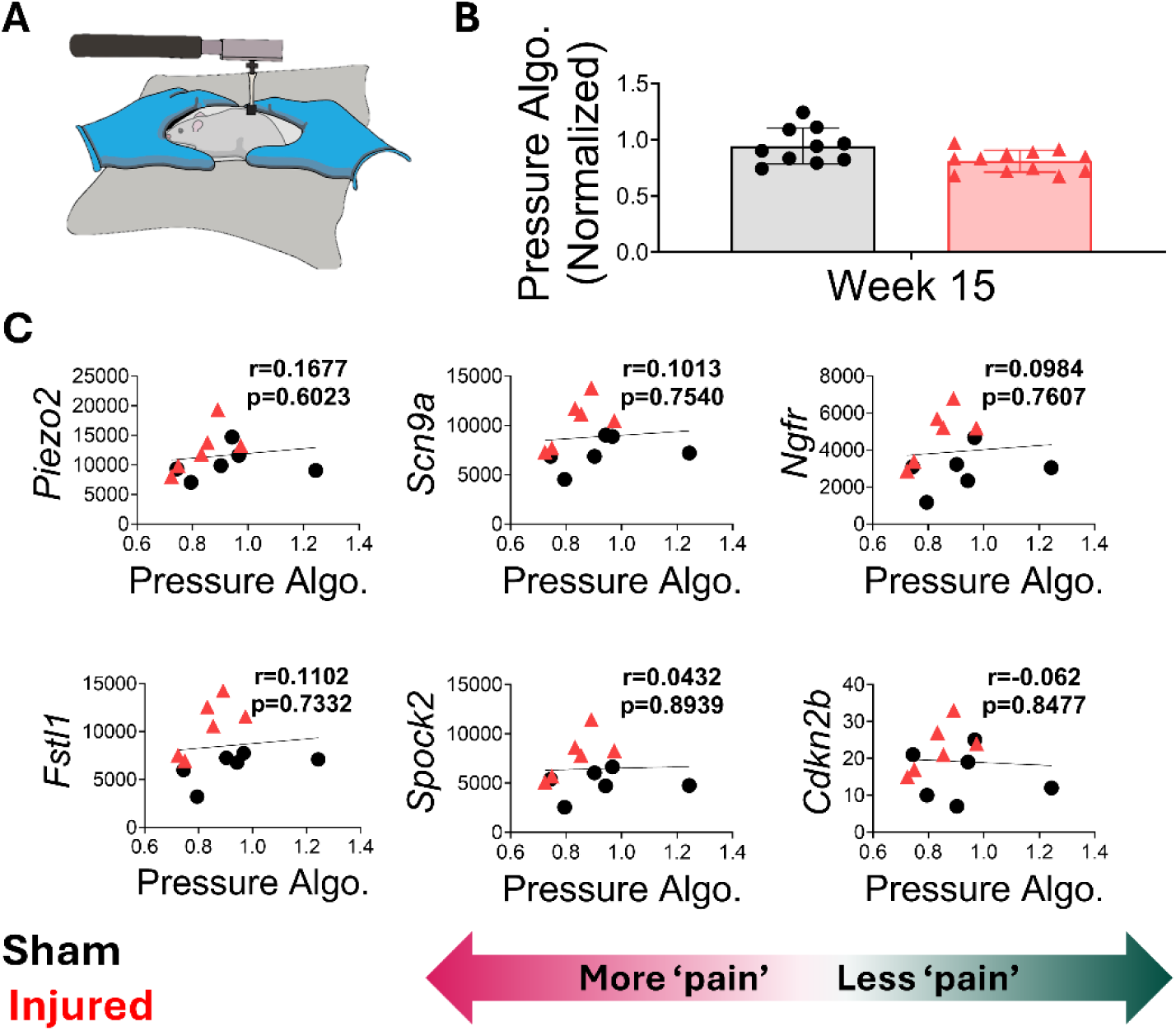
Spearman correlations of differentially expressed genes of interest between sham and injured animals with pain-like behavior measured by pressure algometry. **(A)** Schematic of the pressure algometry assay to evaluate pain-like behavior. **(B)** Normalized pressure algometry at week 15 post-injury, showing significantly lower pressure algometry in injured animals indicating pain-like behavior. **(C)** Spearman correlations between differentially expressed genes relevant to pain-like behavior and disc degeneration (*Scn9a, Ngfr, Fslt1, Spock2, and Cdkn2b*) and pressure algometry. No significance was observed, suggesting that gene expression changes does not correlate with pain-like behavior as measured by pressure algometry in this dataset. N=6 animals per group. Black symbols represent sham, red symbols represent injured animals. *=p ≤ 0.05. The arrow indicates that higher pressure algometry values correspond to less pain-like behavior, reflecting a value closer to the baseline (1).

#### 3.2.3. Differentially expressed pain-related genes do not correlate with CGRP nerves in the disc

Injured discs showed a significant increase in CGRP+ nerve staining compared to sham discs, indicating heightened peptidergic nociceptor innervation post-injury (**Fig. 7A-B**). Specifically, the dorsal annulus fibrosus (AF) exhibited a 5.96-fold increase, the nucleus pulposus (NP) showed a 2.08-fold increase, and the ventral AF demonstrated a 2.79-fold increase in CGRP+ staining (**Fig. 8B**). However, no significant correlations were observed between CGRP staining across the disc and the differentially expressed genes across all clusters (**Fig. 8C**), suggesting that while nociceptor density increases with injury, gene expression profiles in the DRG may not be associated with nociceptive nerve density in the disc.

**Figure 8.**
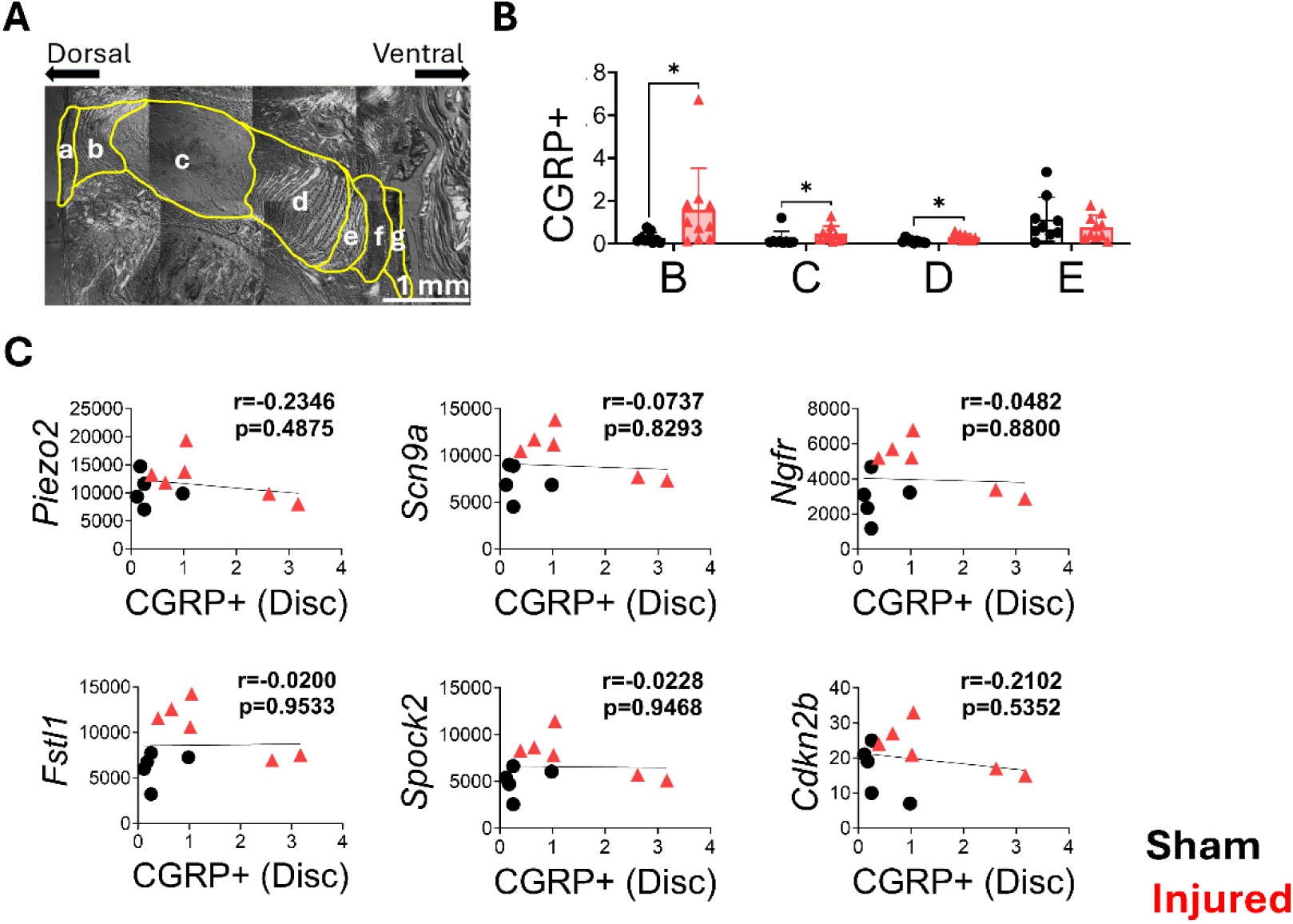
Spearman correlations of differentially expressed genes of interest between sham and injured animals relative to CGRP-positive nerve staining in the disc. **(A)** Diagram of a sham disc segmented into regions of interest for analysis, labeled as follows: a = dorsal ligament, b = dorsal annulus fibrosus (AF), c = nucleus pulposus (NP), d = ventral annulus fibrosus inner 2/3, e = ventral annulus fibrosus outer 1/3, f = granulation, g = ventral ligament. Dorsal and ventral directions of the disc are shown. **(B)** Quantification of CGRP-positive area in the disc at week 15, segmented by disc region. Data show significantly increased CGRP+ staining in the dorsal AF, NP, and ventral AF regions in injured discs, indicating increased peptidergic nociceptors are present post-injury. **(C)** Spearman correlations between pain- and degeneration-related genes (*Scn9a, Ngfr, Fstl1, Spock2, Cdkn2b*) and CGRP-positive area revealed no significant correlations, suggesting gene expression changes do not align with increased CGRP-positive innervation in this dataset. N=6 animals per group. Black symbols represent sham, red symbols represent injured animals. Scale bar = 1 mm.

## 4. Discussion

Discogenic LBP is a complex and persistent condition with limited clinically effective long-term treatments. A key feature of this condition is the abnormal growth of nociceptor axons into previously aneural disc regions [7,20]. Repeated stimulation of nociceptors by noxious mechanical and inflammatory stimuli can elicit plastic changes, as observed in various pain states [26]. These plastic changes include increased expression and presentation of ion channels, enhanced excitability, and a lowered activation threshold [19,26,29]. Despite the prevalence of discogenic LBP, the lack of effective treatments could potentially be attributed to an incomplete understanding of the underlying mechanisms. This study addresses this gap by generating the first comprehensive transcriptional map of DRG neurons in a rat model of discogenic LBP using snRNAseq. These data highlight correlations between changes in nociceptor gene expression, pain-like behaviors, and nerve innervation in the disc. Importantly, these findings reveal significant upregulation of key pain-sensing ion channels, neuroinflammatory markers, and disc degeneration markers in DRGs of LBP animals, suggesting that mechanical hypersensitivity and inflammatory signaling are central to the pain phenotype in this model. Building on prior literature, the following sections will explore specific genes of interest, such as *Scn9a*, *Piezo2*, *Fstl1*, *Ngfr*, and *Spock2* and their potential role in this discogenic LBP model. Additionally, the implications of specific sensory neuron subtypes will be addressed, along with the limitations of this study and directions for future research.

In this study, *Scn9a*, which encodes the Nav1.7 sodium channel, was significantly upregulated in DRGs of injured animals (p-adj≤0.05) across all clusters and significantly correlated with reduced grip strength, a measure of axial hypersensitivity (p=0.0102). This suggests that *Scn9a* may play a role in pain-like behavior in this discogenic associated LBP model. Previous research has demonstrated that upregulation of *Scn9a* lowers activation thresholds in nociceptors, facilitating pain transmission in various chronic conditions, including neuropathic pain and OA [17]. In both an experimental model of neuropathic pain and in human neuropathic patients, DRG *Scn9a* upregulation was linked to enhanced pain sensitivity [38], highlighting its relevance across multiple pain conditions. Taken together, these data highlight the key role that *Scn9a* may play in maintaining hypersensitivity across pain states and specifically in discogenic LBP.

Our findings also demonstrated that *Piezo2*, a key neuronal mechanosensitive ion channel [14,66,69], was significantly upregulated in DRGs of injured animals (p-adj≤0.05) across all clusters. This suggests a component of mechanical hypersensitivity in this model of discogenic LBP. However, *Piezo2* did not significantly correlate with axial hypersensitivity as measured by grip strength (p=0.1267), indicating it may not directly drive the observed pain-like behaviors in this model. Prior studies have shown that *Piezo2* transduces mechanical signals into neuronal activation and contributes to pain sensitivity, with a well-documented role in mechanical allodynia in preclinical pain models [51,54]. These prior studies suggest the role of *Piezo2* as key driver in conditions characterized by mechanical pain, where nociceptor excitability is increased in response to mechanical stimuli. *Piezo2* has an established role in various pain models [51,54], and showed a log2 fold change of 0.44 in our discogenic LBP model. Although no correlations were observed between overall *Piezo2* expression and pain-like behaviors in this model, *Piezo2* expression was increased in pain-sensing nerve subclasses, particularly in Ntrk3low+Ntrk2 Aδ-LTMRs. Therefore, *Piezo2* may remain a target for further examination in discogenic LBP models.

Our findings demonstrated significant upregulation of *Fstl1* (Follistatin-like protein 1) (p-adj≤0.05) and *Ngfr* (p75NTR) (p-adj≤0.05) in DRGs of injured animals across all clusters, suggesting a role for neuroinflammation in this model. Notably, *Fstl1* expression strongly correlated with axial hypersensitivity (p=0.0025), suggesting its potential contribution to pain-like behavior in this model. *Fstl1* has been shown to be elevated in inflamed and degenerative disc tissues in a rabbit model of intervertebral disc degeneration [75]. *Fstl1* promotes inflammation and nucleus pulposus cell senescence through the TLR4/NF-κB pathway, exacerbating disc degeneration [75], which suggests a key role in chronic LBP. Additionally, *Ngfr*, which encodes the p75 nerve growth factor receptor, plays a critical role in neurotrophic signaling and nociceptor plasticity [67]. NGF signaling through p75NTR and TrkA receptors has been shown to support nerve innervation and promote nociceptor sensitization in animal models of OA [3]. Increased *Ngfr* expression in our injured animals across all clusters suggests that neuroinflammation may also contribute to the persistent pain observed in our animals. These data highlight the potential role of inflammatory signaling in sustained nociceptor plasticity across various chronic pain states. Our findings revealed a significant upregulation of *Spock2* (p≤0.05), a gene associated with extracellular matrix integrity and remodeling, across all clusters in injured DRGs of this discogenic LBP model (**Fig. 4D**). Further, *Spock2* was significantly correlated with axial hypersensitivity in this model (**Fig. 5C**) (p=0.0205). In this model, the increased expression of *Spock2* may reflect active disc degeneration and its effects on nociceptor plasticity within the DRGs. *Spock2* has been identified as a potential biomarker for LBP patients. Notably, genome-wide association studies from the UK Biobank identified *Spock2* as a genetic risk factor for low back pain, linking this gene to the pathophysiology of disc degeneration and pain in human patients [21]. Together, these findings suggest *Spock2* may serve as both a biomarker of disc degeneration and discogenic LBP. This study highlights DRG subtype-specific changes in Ntrk3low+Ntrk2 Aδ-LTMRs and Calca+Sstr2 peptidergic C-fibers, which exhibited increased proportions in injured animals (3.3% and 2.3% expression in injured and sham animals, respectively). Ntrk3low+Ntrk2 Aδ-LTMRs are associated with hair follicle innervation and detect stimuli on the skin through Ntrk2 (brain derived neurotrophic factor receptor) and Ntrk3 (NT-3 receptor) [1]. A study by Dhandapani et al. [16], showed that mice lacking this Aδ-LTMRs of Ntrk2-positive neurons exhibit reduced touch sensitivity and fail to respond to mechanical stimulation after neuropathic injury, suggesting that these Aδ-fibers may contribute to the development of allodynia. These fibers showed significant upregulation of pain-relevant genes such as *Scn9a*, *Piezo2*, and *Ngfr*, implicating these genes in heightened sensitivity observed in this chronic LBP model. Although their presence in painful intervertebral discs is unknown, their involvement in allodynia suggests a potential role in discogenic pain. Similarly, Calca+Sstr2 C-fibers (8.5% and 7.1% expression in injured and sham animals, respectively), showed elevated expression of *Spock2* and *Fstl1*, suggesting these subtypes play a key role in driving pain and structural degeneration in discogenic LBP. In the Calca+Smr2 C-fiber population, genes such as *Ngfr*, *Calm1*, *Kcna2*, and *Calca* were differentially expressed in injured animals, suggesting these genes may have a role nociceptor plasticity in this DRG neuron subtype. Notably, *Calca*, which encodes CGRP—a key mediator of pain and inflammation [72]— showed increased expression. This was further supported by elevated CGRP+ nerve staining in the disc, indicating heightened nociceptor innervation post-injury, particularly within the dorsal annulus fibrosus.

There are some limitations to our study. First, our findings are based on a rat model. Although our LBP model recapitulates many aspects of human LBP, it does not fully capture the complexity of human DRG structures or nociceptor diversity [5,64]. Importantly, when annotating the DRG dataset for this study it was observed that the rat DRGs were more closely related to mice than humans. During analysis, 18 neuronal subtypes were identified across mouse and rat DRG datasets, using Conos label transfer. This study highlights both conserved and species-specific transcriptional profiles. Notably, Ntrk3high+Ntrk2 Aβ-RA LTMRs were more prevalent in rats, while Th C-LTMRs were more prevalent in mice, suggesting interspecies differences in sensory neuron subpopulations. The rat dataset revealed 49 unique genes absent in mice, including Dsp (cell adhesion and structural integrity) and Nrdc (regulation of growth factors and inflammatory response), which may contribute to species-specific injury and repair mechanisms. These differences underscore the need to consider species-specific neuronal distributions and gene expression patterns in pain studies.

An additional limitation is that analyzed gene expression was analyzed at a single time point (15 weeks post-injury), capturing chronic stage molecular profiles but potentially missing earlier dynamic changes in DRG plasticity. Longitudinal studies capturing gene expression at multiple time points would provide insights into the progression of nociceptor changes and could reveal potential intervention points [76]. In our study, a small percentage of *Atf3* cells (0.13%) were observed. Studies of various pain models have examined significant changes in *Atf3*, a marker of nerve injury, in timepoints up to 28 days [9,46]. However, due to our extended timepoint of 15 weeks post-injury, the transient expression changes in *Atf3* or other acute markers of neuronal injury and stress may be missed. Lastly, no interventional studies were conducted to examine the potential role of genes that showed altered gene expression in pain in this LBP model. This can be assessed in future studies, in particular for Nav1.7. Along those lines, the findings highlight the potential of DRG-targeted interventions. Another opportunity would be a nociceptor-specific *Piezo2* knockout model like that used in experimental OA demonstrated significant reductions in pain-like behaviors and joint damage [54]. These results suggest that similar strategies could be adapted for discogenic LBP, with a focus on targeting DRG-specific mechanisms to alleviate pain and improve clinical outcomes. Since these models are more amenable to using rats, genetic tools for targeting sensory neurons in this species would be useful.

To conclude, this study provides insights into the molecular changes driving chronic discogenic LBP by offering the first comprehensive transcriptional map of DRG neurons using snRNAseq in a rat model. This work establishes a foundational rat atlas of DRG subpopulations, identifying 18 unique clusters and emphasizing the importance of nociceptor subtype-specific changes in discogenic LBP. Key genes, including *Scn9a*, *Ngfr*, *Fstl1*, and *Spock2*, were significantly correlated with axial hypersensitivity. This work establishes a foundation for understanding the molecular mechanisms underlying chronic discogenic LBP, and highlights opportunities for developing targeted, non-opioid therapies to enhance therapeutic translation and patient outcomes.

## Acknowledgments

The authors thank Kayla Ney and Evie Barnett for their support with pressure algometry and disc histology and immunohistochemistry, respectively. The authors thank the Genome Core at the University of Texas at Dallas for technical support. This work was funded by the National Institutes of Health (R01AR080926) and supported by the NIH grant (NS065926) to the Center for Advanced Pain Studies at the University of Texas at Dallas. Sequencing data is available at GEO at accession number GSE289659.

